# Evaluation of multi-peptide sequences identified *in silico* for the serological identification of antibodies against Vector-Borne Diseases

**DOI:** 10.1101/2024.04.16.589708

**Authors:** Carlos A. Peña-Bates, Cesar I. Lugo-Caballero, Norma Pavía-Ruz, Henry R. Noh-Pech, Oghenekaro Omodior, Fernando I. Puerto-Manzano, Karla R. Dzul-Rosado

## Abstract

The socio-ecological conditions of Mexican regions are conducive to the spread of vector-borne diseases. Although there are established treatment guidelines for dengue and rickettsiosis, diagnosis is complicated. The objective of this work was to identify epitopes of *Rickettsia* and Dengue virus that could be used in serology screening against vector-borne diseases. For this, epitopes with high HLA-II binding efficiency of OmpB protein of *Rickettsia rickettsii* and NS2B protein of dengue virus were identified *in silico* through a reverse vaccinology strategy. The selected epitopes were grouped into multi-peptide sequences that were synthesized and immobilized in a nitrocellulose membrane to evaluate the reactivity sera from patients previously infected with dengue or *Rickettsia*. The evaluation of the sequences of the NS2B and OmpB proteins was performed with 60 sera previously diagnosed as positive or negative by the respective gold standard techniques. The *dot blot* technique was used for the antigenic evaluation of the peptides against these serum samples. *Dot blot* analysis correctly identified 85% of sera positive for rickettsiosis and 75% of sera positive for dengue. Reverse vaccinology reduces the time and cost of antigen discovery. Experimental evidence from multi-peptide sequences suggests their potential use in the development of diagnostic tests for dengue and rickettsiosis.

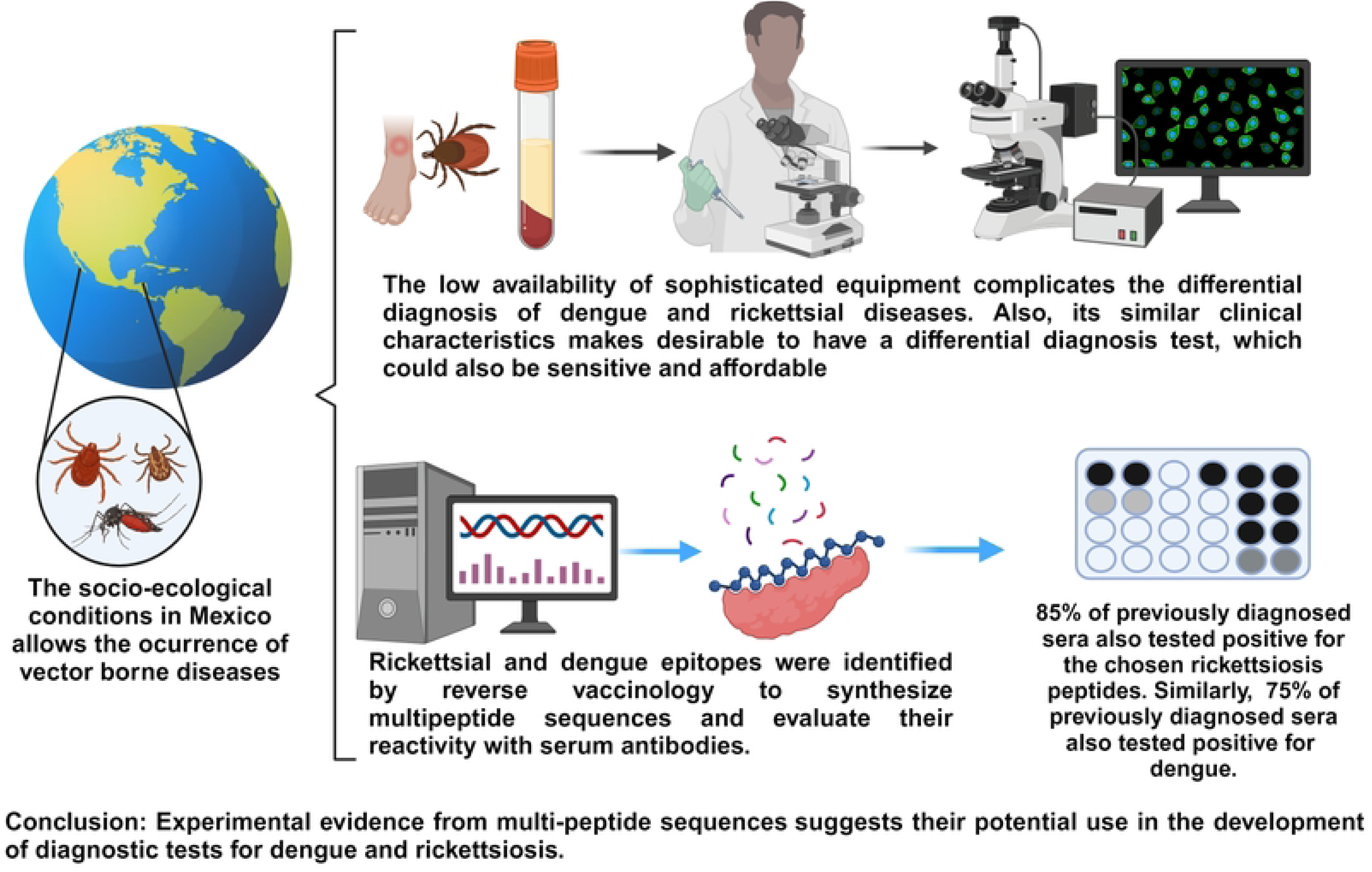

## 1. INTRODUCTION

Vector-borne diseases (VBDs) account for about 17% of infectious diseases worldwide.^1^ Among these diseases, some are viral, such as dengue, zika, or chikungunya, transmitted by the bite of mosquitoes of the genus *Aedes* (species *aegypti* and *albopictus*)^2^ and some are bacteria, such as rickettsiosis, transmitted mainly by the bite of fleas and ticks.^3^

The nonspecific clinical symptomatology in the first days of the febrile phase, characterized by myalgia, arthralgia, fever, and rash, makes difficult the diagnosis of dengue, rickettsiosis, and other diseases that exhibit similar clinical characteristics (chikungunya, zika, etc.). In addition, the difficulty of access to diagnostic tests and the low interest in performing confirmatory laboratory tests lead to late treatment of patients with a misdiagnosis, with fatal consequences in some cases.^2,4^

Laboratory diagnosis of dengue and rickettsial diseases is based on molecular and serologic tests, the latter being the most widely used. The WHO recommends the use of the IgM antibody capture ELISA technique for the diagnosis of dengue, which remains the gold standard for the diagnosis of the disease.^5,6^

In the case of rickettsiosis, Indirect Immunofluorescence (IFI) is the serological test accepted as the gold standard for diagnosis in Mexico and the world; however, it has the disadvantages of requiring specialized equipment, low sensitivity, and specificity due to the time for the presence of the first antibodies (7-10 days) and to the fact that the commercial kits are limited to a few species of *Rickettsia*.^7,8^

It is important to mention that the main difficulty of diagnostic tests for dengue in the clinical laboratory is the high cross-reactivity with other *Flaviviruses* (Zika virus, West Nile virus, and Yellow Fever virus), which does not allow for accurate serologic diagnosis. On the other hand, the diagnosis of rickettsial diseases presents several challenges, mainly due to the cost of IFI tests and the limited availability of the necessary equipment and infrastructure reducing its availability. ^8,9^

Due to the epidemiological reemergence of these diseases, the development of a serological diagnostic system based on synthetic peptides could be operationally cheaper, faster, easier to produce, and capable of diagnosing dengue or rickettsiosis with relevant sensitivity and specificity however, the identification of a good antigen may be a slow process.^10^

A novel and faster option for peptide selection is through *in silico* identification, by predicting specific epitopes toward the major histocompatibility complex class II (HLA-II) haplotypes circulating in our region and analyzing them with software that predicts their presence and affinity in CD4+ T-cells and B-cells. This technique, known as reverse vaccinology, leads to the expedited recognition of antigens. These antigens are sorted by their antigenicity and then assessed for their reactivity via experimental methods. The potential cross-reactivity with other microorganisms must be validated and evaluated in a subsequent stage.^11–13^

This study employs a bioinformatic prediction-based (*in silico*) strategy that aims to identify immunoreactive epitopes located within conserved antigenic regions of *Rickettsia* and Dengue virus. Subsequently, they are evaluated through serological analysis to determine their usefulness as candidate antigens for validation in a rapid diagnostic test.

## 2. MATERIALS AND METHODS

### 2.1. Serological samples

Serum samples were obtained from the Serum Bank of the Laboratory of Emerging and Re-emerging Diseases and the Hematology Laboratory of the Regional Research Center Dr. Hideyo Noguchi of the Autonomous University of Yucatan All samples collected between February and December 2014 were obtained by venipuncture, and informed consent was obtained for this study.

Serum samples were previously tested by ELISA (PanBio Dengue duo IgM and IgG) and immunochromatographic antibody test (Dengue combo Ab (IgG+ IgM) + Ag (NS1)) against Dengue virus and IFI for *Rickettsia rickettsii* and *Rickettsia typhi*. Twenty (20) IgG sera positive for Dengue virus, 20 IgM/IgG sera positive for *Rickettsia*, and 20 IgM/IgG sera negative for both pathogens were selected.

### 2.2. Ethical considerations

This study was reviewed and approved by the Research Ethics Committee of the Regional Research Center “Dr. Hideyo Noguchi” of the Autonomous University of Yucatan (CIRB-2015-005). Prior to serological sample collection, each study participant provided a signed informed consent sheet.

### 2.3. Protein identification

The amino acid sequence of the full-length polyprotein of the dengue virus type 2 (DENV2) was retrieved from Genbank with accession number AHI43693.1. The amino acid sequence of the full-length polyprotein of the dengue virus type 2 (DENV2) was retrieved from Genbank with accession number AHI43693.1. The BLASTp platform was used to identify individual protein sequences similar (>90%) to the viruses: Zika, Chikungunya, and West Nile, which were discarded. The NS2B protein was selected for its degree of conservation among the four serotypes of Dengue virus (≥ 80%); besides, in this protein, there is a region of 60 amino acids that is not conserved with Zika virus, others *Flaviviruses* and *Rickettsia sp*.

Based on vaccine and diagnostic test development studies,^14–16^ the OmpB protein of *Rickettsia rickettsii* (WP_012151219.1) was used and compared to determine its similarity and intraspecies conservation with seven human pathogenic species: *Rickettsia felis, Rickettsia akari, Rickettsia typhi, Rickettsia prowazekii, Rickettsia slovaca, Rickettsia conorii,* and *Rickettsia parkeri*, which were selected based on their identification in reported human cases of rickettsioses in Mexico and the Americas. A second BLASTp analysis was performed with other organisms (*Leptospira sp*, Dengue virus, Zika virus, and Chikungunya virus) to identify and exclude similar protein regions of other pathogens.

### 2.4. Epitope prediction and peptide selection

Using the 970-1260 amino acid sequence of the OmpB protein and the 60 amino acid region of the NS2B protein, prediction programs were used to identify immunogenic epitopes. HLA class II molecules present in TCD4+ cells were predicted using the NetMHCIIpan 3.2 and NetMHCII2.3 servers, accessible at http://www.cbs.dtu.dk, with cutoffs of 0.60 and above. The HLA class II haplotypes with the highest frequency in the Mexican population (HLA-DRB1*01:01, HLA-DRB1*03:01, HLA-DRB1*04:01, HLA-DRB1*04:02, HLA-DRB1*04:04, HLA-DRB1*04:05, HLA-DRB1*07:01, HLA-DRB1*08:02, HLA-DRB1*09:01, HLA-DRB1*11:01, HLA-DRB1*13:02, HLA-DRB1*15:01) were included in the analysis.^17,18^ These haplotypes were selected according to data from the Allele Frequency Net Database (AFND), based on their percentage frequency in Mexico. Predictions were made with linear peptides of variable length from 12 to 15 amino acids.

The selected epitopes’ potential presentation in HLA class-II molecules of B-cells were evaluated through several algorithms to inquire their epitope specificity (BEPIPRED) (score of 0.50-0.55), flexibility and torsion angle (Karplus and Schulz) (score ≥ 0.90); hydrophilicity index (Parker) (positive score); antigenicity scale (Kolaskar and Tongaonkar) (score ≥ 1.0) and surface accessibility for linear epitopes (Emini) (score ≥ 0.5). These were evaluated Using the Immune Epitope Database and Analysis Resource (IEDB) tool available at: http://www.iedb.org/. The peptides were assessed with the ABCpred software to enhance the reliability of the epitope selection strategy.

Finally, the selected epitopes were arranged into a multi-peptide sequence using residue alignment and overlapping, following the procedure described by Mishra *et al*.^19^ A new alignment was performed using the MultAlin program (available at: http://www.sacs.ucsf.edu/cgi-bin/multalin.py) to determine the degree of sequence conservation in the dengue virus serotypes greater than 65% and species selected in the case of *Rickettsia* sp greater than 95%.

### 2.5. Peptide synthesis

The multi-peptide sequences were synthesized in Peptide2.0 laboratories (USA) with a purity greater than 80%.

### 2.6. Dot Blot

Using the Bio-Dot device (BIO-RAD), nitrocellulose membranes (BIO-RAD) were sensitized for each multi-peptide sequence by applying 1.25 μg/dot (50 μL) of the synthetic peptide as antigen dissolved in PBS1X/BSA1% for 2 hours. The membrane was blocked with PBStween/BSA1%. The membrane was then incubated with control sera (negative and positive, and samples for each pathogen) at serial dilutions from 1:1,000 to 1:10,000. A wash was performed with 100 μL PBS/Tween for 30 minutes. Peroxidase-linked anti-IgG or human IgM conjugate secondary antibodies (IgM: Thermo Scientific, USA #31415; IgG: PROMEGA, USA #W4038) were added at a 1:5,000 dilution with PBStween/BSA for 1 hour. After a 30-minute wash with PBS/Tween, the detection was performed with 4-chloronaphthol (Sigma Aldrich, USA #C8302) with 30% hydrogen peroxide. A contact time of three minutes with the developing solution was set. The reaction was stopped by washing the membrane with doubly distilled water and the result was observed without allowing it to dry or for more than five minutes.

## 3. RESULTS

### 3.1 Prediction and Selection of Peptides

The selected proteins have to be specific for the microorganism, so discarding the homologous proteins with other organisms is necessary. In the case of the Dengue virus, the candidate was the NS2B protein, presenting a similarity of ≥ 65% between the four serotypes of the virus, including a zone of 60 amino acids not shared with other *Flaviviruses*. In the case of *Rickettsia sp*, the OmpB protein was selected, which has a degree of similarity ≥ 95% between the species of the genus included.

The NS2B and OmpB proteins were analyzed with NetMHCII and NetMHCIIpan servers, which identify by prediction epitopes of binding to MHC class II molecules of CD4+ T-cells that are important for activation and differentiation of B-cells.

For the NS2B protein, 1,320 epitopes were predicted, of which 91 were present in both NET prediction programs, and 31 promiscuous epitopes were selected for different HLA class-II haplotypes: HLADRB1-0101, -0701, -0901, -1302 (Figure 1).

**Figure 1.**
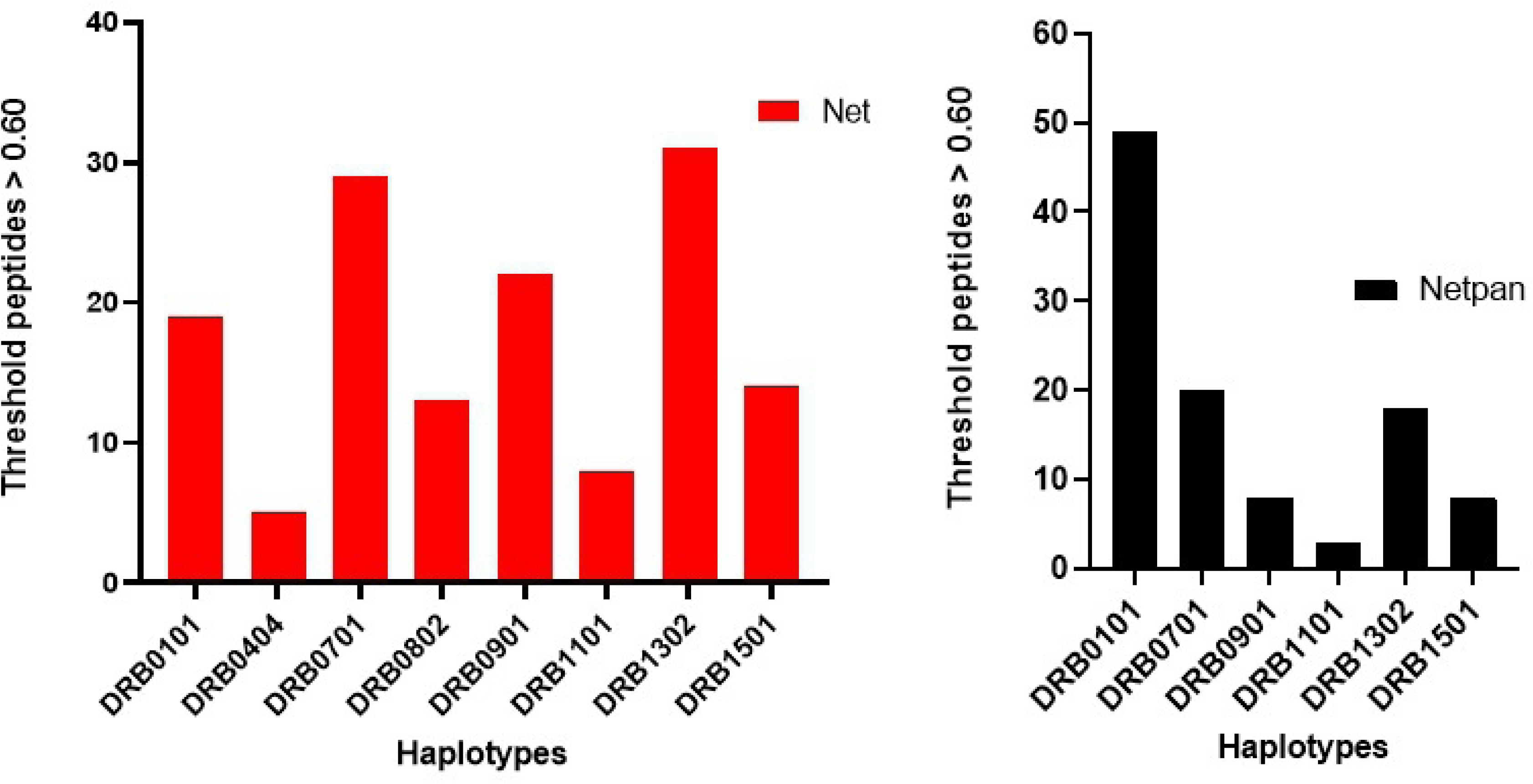
Graphs corresponding to peptides with binding affinity to HLA class-II haplotypes for the NS2B protein of DENV2. On the left is the chart comparing to the NetMHCII 2.3 server program of experiments with mice. On the right side, the graph corresponds to the experimental NetMHCIIpan program of humans.

For the OmpB protein, 23,200 epitopes were predicted, of which 453 were present in both NET prediction programs. Of the 453, they turned out to be 180 peptides promiscuous to different HLA class-II haplotypes HLADRB1-0101, -0701, -0901, -1101, and -1302 (Figure 2).

**Figure 2.**
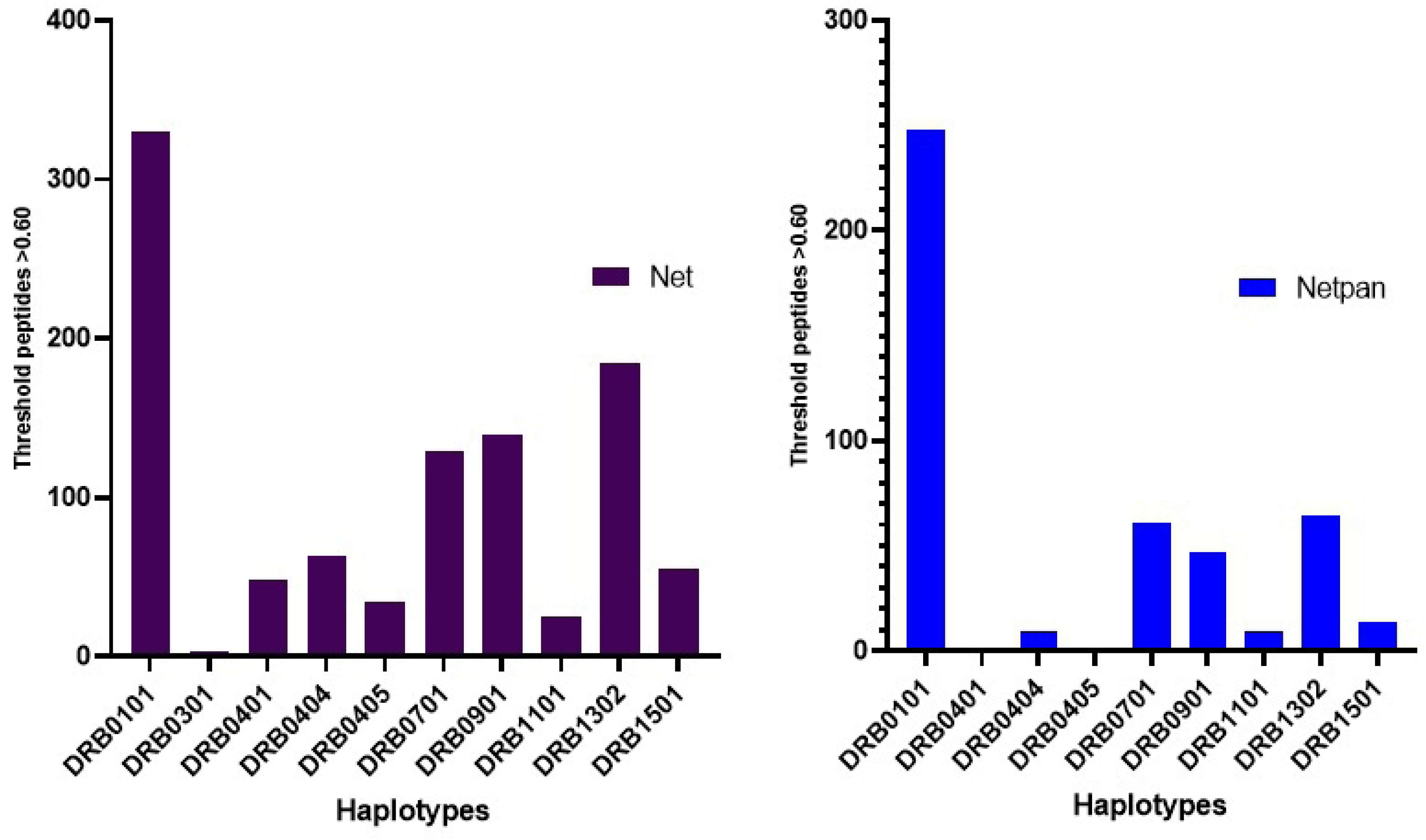
Graphs corresponding to peptides with binding affinity to HLA class-II haplotypes for Rickettsia protein OmpB. On the left is the chart comparing to the NetMHCII 2.3 server program of experiments with mice. On the right side, the graph corresponds to the experimental NetMHCIIpan program of humans.

The selected epitopes were evaluated using the IEDB and ABCpred analysis resource to predict different physicochemical properties and select those viable peptides to be presented by B lymphocytes to TCD4 + cells. In the end, seven epitopes of the NS2B protein were selected (Table 1), and nine epitopes of the OmpB protein with high antigenicity scores and outstanding physicochemical characteristics (Table 2).

**Table 1.** Antigenic epitopes of Dengue virus type 2 (DENV2) towards selected CD4+ T-cells and their values obtained in the physicochemical properties in B-cells.

| Peptides | Size | Nucleus | Average Nets | B | E | K | P | KA |
| --- | --- | --- | --- | --- | --- | --- | --- | --- |
| TGLLVISGVFPVSIP | 15 | LVISGVFPV | 0.65 | 0.549 | 0.20 | 1.12 | -1.39 | 0.96 |
| FPVSIPITAAAWYL | 14 | SIPITAAAW | 0.62 | 0.521 | 0.55 | 1.10 | -1.99 | 0.94 |
| GVFPVSIPITAAAW | 14 | FPVSIPITA | 0.62 | 0.524 | 0.31 | 1.08 | -1.05 | 0.96 |
| VSIPITAAAWYLWE | 14 | PITAAAWYL | 0.61 | 0.522 | 0.75 | 1.07 | -1.64 | 0.92 |
| FPVSIPITAAAWYLW | 15 | SIPITAAAW | 0.61 | 0.521 | 0.61 | 1.08 | -2.52 | 0.94 |
| SGVFPVSIPITAAA | 14 | FPVSIPITA | 0.62 | 0.526 | 0.40 | 1.09 | 0.13 | 0.96 |
| TGLLVISGVFPVSI | 14 | LVISGVFPV | 0.69 | 0.551 | 0.12 | 1.13 | -1.64 | 0.96 |
| Peptides with the best scores in ABCpred | TILIRTGLLVISGVFP (score:0.94), SIPITAAAWYLWEVKK (score:0.92), LVISGVFPVSIPIT (score:0.84), VSIPITAAAWYLWE (score:0.80) |  |  |  |  |  |  |  |
| B: BEIPRED, E: EMINI, K: KOLASCAR, P: PARKER, KA: KARPLUS |  |  |  |  |  |  |  |  |

**Table 2.** Antigenic epitopes of the OmpB protein of Rickettsia towards selected CD4+ T-cells and their values obtained in the physicochemical properties in B-cells.

| Peptides | Size | Nucleus | Average Nets | B | E | K | P | KA |
| --- | --- | --- | --- | --- | --- | --- | --- | --- |
| SPNFAVTGSNRFV | 13 | FAVTGSNRF | 0.81 | 0.554 | 0.71 | 1.024 | 1.577 | 1.021 |
| SPNFAVTGSNRFVNY | 15 | FAVTGSNRF | 0.80 | 0.550 | 1.24 | 1.02 | 1.71 | 1.03 |
| PNFAVTGSNRFVNY | 14 | FAVTGSNRF | 0.80 | 0.550 | 1.10 | 1.02 | 1.36 | 1.04 |
| PNFAVTGSNRFVNYS | 15 | FAVTGSNRF | 0.79 | 0.548 | 1.24 | 1.02 | 1.71 | 1.03 |
| NFAVTGSNRFVNY | 13 | FAVTGSNRF | 0.79 | 0.550 | 0.864 | 1.013 | 1.308 | 1.05 |
| PNFAVTGSNRFVN | 13 | FAVTGSNRF | 0.79 | 0.552 | 0.852 | 1.006 | 1.615 | 1.038 |
| NRFVNYSLIRAAQ | 14 | YSLIRAAQ | 0.72 | 0.529 | 1.53 | 1.02 | 1.01 | 0.94 |
| RFVNYSLIRAAQD | 14 | YSLIRAAQ | 0.72 | 0.528 | 1.59 | 1.03 | 1.22 | 0.94 |
| RFVNYSLIRAAQ | 13 | YSLIRAAQ | 0.71 | 0.528 | 1.16 | 1.04 | 0.55 | 0.93 |
| **Peptides with the best scores in ABCpred** | NRFVNYSLIRAAQ (score:0.75), SPNFAVTGSNRFVN (score:0.53), NFAVTGSNRFVNYSLI (score: 0.78) |  |  |  |  |  |  |  |

The NS2B protein sequence consists of 26 amino acids: **TGLLVISGVFPVSIPITAAAWYLWE**, which is located between residues 1446-1470 of the Dengue virus type 2 sequence. Through multiple alignments, the consensus sequence of the NS2B protein reveals its location within a semi-conserved region among the four serotypes of the Dengue virus. However, it lacks highly conserved regions when compared to the Zika virus or West Nile virus. (Figure 3).

**Figure 3.**
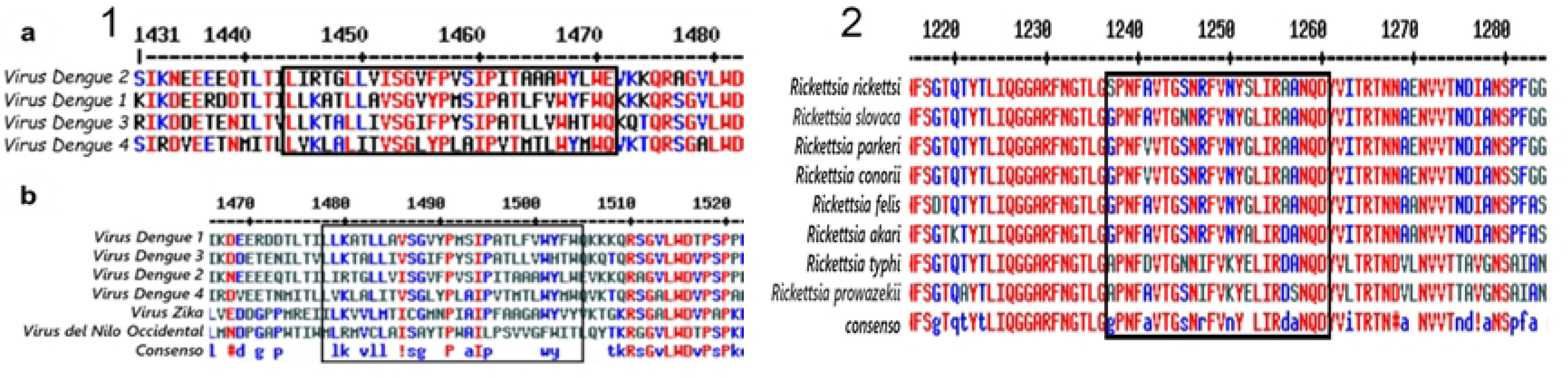
Localization of epitopes in multiple alignments. 1) multiple alignments of the NS2B protein. a) Multiple alignments of the four dengue virus serotypes, b) alignment of the four dengue virus serotypes with Zika virus and West Nile virus, the box indicates the peptide sequence synthesized. 2) multiple alignments of Rickettsia’s OmpB protein. In the black box, it is located in a semi-conserved region among the species included in the study.

The OmpB protein sequence consists of 24 amino acids: **SPNFAVTGSNRFVNYSLIRAANQD**, which is located between residues 1220 and 1243 of the complete sequence of *Rickettsia rickettsii*. In multiple alignments, it is observed that the sequence is in a region mostly conserved in all *Rickettsia* species included in the study (Figure 3).

### 3.2. Determination of anti-DENV2 IgG antibodies from characterized sera from infected patients

A total of 40 serologic samples previously characterized as positive and negative by ELISA were evaluated, of which the dot blot confirmed that 75% (15/20) were positive for IgG antibodies. It was also found that 80% (16/20) of the sera that were true negative by ELISA were also negative by dot blot using secondary IgG antibodies (Figure 4)(S1).

**Figure 4.**
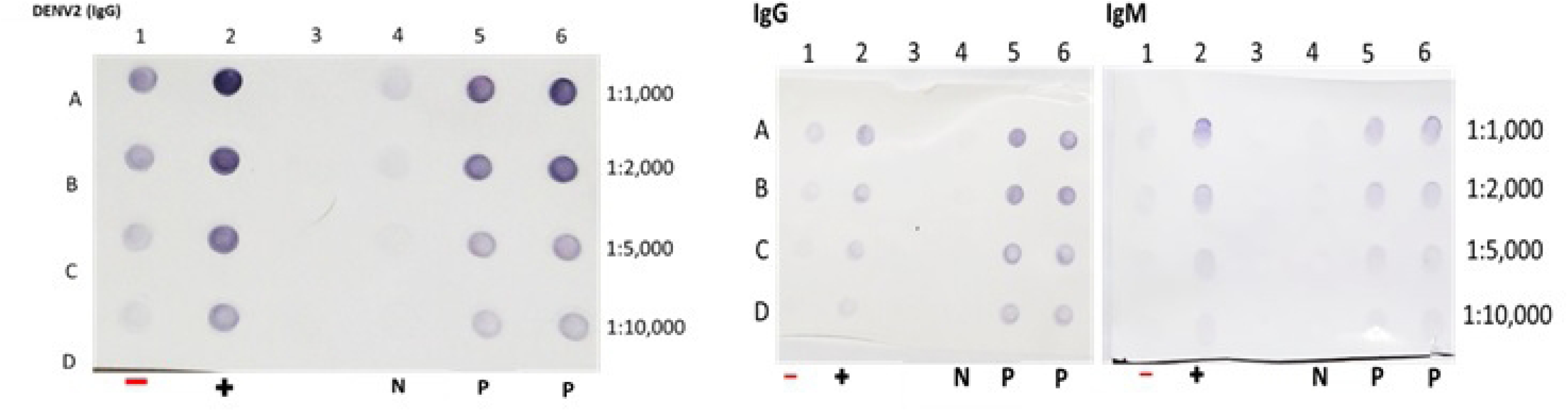
Dot blot representative of the sera evaluated. For the NS2B protein of the DENV2 virus (left). Representative image in the interpretation of sera for IgG antibodies. On the right side, there are two membranes of the Dot blot representative of the sera evaluated for the OmpB protein in the interpretation of the sera for the IgG and IgM antibodies. In all the images we have: lane 1 represents the negative control serum (-), 2 the positive control serum (+), 3 places all the reagents except the problem serum, 4 is a negative sample (N), and lanes 5 and 6 represent positive samples (P).

### 3.3. Determination of anti-Rickettsia IgM/IgG antibodies from characterized sera of infected patients

A total of 40 serological samples previously characterized as positive and negative by Indirect Immunofluorescence (IFI) were evaluated, of which the *Dot blot* confirmed that 85% (17/20) were positive for IgG antibodies and 75% (15/17) for IgM antibodies. In the case of true negative sera, it was confirmed by *Dot blot* that 85% (17/20) was for IgG and 82% (19/23) for IgM, using secondary antibodies IgG or IgM depending on the experiment (Figure 4) (S2).

## 4. DISCUSSION

The increasing spread of vectors due to global warming and migration has made dengue and rickettsiosis a threat to the Mexican population. Access to the health care system which is necessary for the availability of expensive diagnostic equipment and methods is absent in rural communities, leading to late treatment and fatal outcomes.^17,20^

Available diagnostic methods such as IFI and PCR for rickettsiosis or ELISA for dengue have worked adequately, but in many regions of our country, they are unavailable or expensive and require time, materials, and personnel with extensive experience. It has also been observed that the type of *Rickettsia* antigen used in commercial tests originates from microorganisms that are scarce or absent in most parts of Mexico. For dengue, serological and Immunochromatographic tests cross-react with other pathogens such as the Zika and other *Flaviviruses*, and the Chikungunya virus.^6,7,10,21^ Against this background, epidemiologic surveillance of both diseases requires new methods of serologic diagnosis that are rapid, accessible and do not require expensive equipment, thus avoiding late treatment and fatal outcomes.

This study was based on a bioinformatics analysis using prediction programs based on experimental data and evaluated by artificial neural network algorithms. The results obtained for the selected peptides and their experimental analysis show that the reverse vaccinology strategy is efficient in the identification of antigenic peptides with specific binding to antibodies, making it useful for the development of novel and accessible diagnostic methods.^10,12^

In the case of the Dengue virus, the candidate in the analysis of the BLASTp program was the NS2B protein. This transmembranal protein participates as a cofactor with the NS3 protein in viral replication. It has also been described that its antigenic presentation in the generation of antibodies in BALB/c mice is mediated by HLA class II molecules, generating a limited IgG antibody response against it in the course of primary infection and increased when they undergo a second infection of the disease by any of the four serotypes of the virus.^22,23^ On the other hand, the region of interest in this bioinformatics analysis is not conserved with other *Flaviviruses.* The assessment of non-structural proteins that produce IgG antibodies is a beneficial technique to determine dengue exposure in endemic populations and predict the possibility of dengue infection with severe symptoms. Currently, there are no dengue-specific serological tests available due to the high prevalence of cross-reactivity with other *Flaviviruses*. This cross-reactivity is a result of B-cells development against conserved epitopes after exposure to the pathogen.^24^

The surface protein OmpB of *Rickettsia* belonging to the Sca family was used. The Sca5 or OmpB protein is found in almost all groups of the genus *Rickettsia* with a molecular weight of 168 kDa. In general, it acts in conjunction with the OmpA protein, mediating the adhesion and invasion of the host cell.^21,25^ It is also known that the OmpB protein is highly immunogenic and has been studied and used to obtain recombinant proteins from the genome of *R. conorii* where it has obtained optimal values of sensitivity and specificity.^16^

It was critical to determine the length of the peptides to be evaluated because the binding cleft of the HLA class II molecule is open and allows the coupling of peptides ranging in size from 12 to 50 amino acids.^26–28^ However, the variability in peptide size still needs to be properly standardized in prediction programs. Therefore, a length of 12 to 15 amino acids has been used. Since the sequence size defines the peptide core, the flanking amino acids determine the affinity and immunogenicity of the peptide, which may result in low accuracy compared to predictions made for HLA class I molecules limited to nine amino acids. In this study, short peptides were not evaluated experimentally. Nevertheless, they were aligned to form a multi-peptide sequence, taking into account the results obtained by Mishra *et al*.,^19^ since they demonstrated that short peptides tend to have low reactivity, less than 60% in the antigen-antibody evaluation.

Regarding the results obtained in the evaluation of antigen-antibody reactivity for the peptide TGLLVISGVFPVSIPITAAAWYLWE of Dengue virus type 2 (DENV2), reactivity was observed in 15/20 sera positive for IgG antibodies, with five results obtained as false negatives. It was also noted that four samples tested gave a false positive effect. Regarding the results obtained for the peptide SPNFAVTGSNRFVNYSLIRAANQD of *Rickettsia* derived from the OmpB protein, reactivity was observed in 17/20 sera positive for IgG and 15/17 sera positive for IgM. Three samples tested gave a false negative result for IgG and another two gave a false negative result for IgM.

The false positives and negative results obtained with some sera could be related to the characteristics of the amino acids that make up the selected peptides by influencing the formation of the peptide core and determining the flanking sites, a fundamental part of the immunogenicity associated with the use of synthetic peptides.^26,27^ Other factors that could influence the interpretation and obtaining of the results would be errors in the understanding of the result, being a qualitative and visual technique, so it depends on the appreciation of the analyst researcher and the inherent properties of nitrocellulose membranes that interact with some characteristics of the amino acids present in the peptide, promoting a low binding of the peptide or blocking essential target sites for the correct interaction of the peptide with the antibodies present in the sample^26,29^, therefore other types of membranes such as PVDF or Z should be tested. Another method that can be used is quantification by optical densitometry, we use histogram evaluation with the ImageJ program, but it is not in the interest of the project to evaluate relative expression levels or to quantify antigen-antibody binding, for which it would be convenient to migrate to a more robust technique such as ELISA.

The results obtained with the peptides suggest that they may be useful in the evaluation of patients with suspected clinical symptoms of dengue or rickettsiosis; however, there are no data available to compare with other diagnostic tests using synthetic peptides of the proteins used. In support of this hypothesis, the study carried out by Mishra *et al.*^19^ Was for Zika virus and used the NS2B protein in a region close to that found in this study, suggesting that this region may be helpful in the differential diagnosis of diseases caused by Dengue virus and Zika virus. This will need to be evaluated in another study.

As this is the first study using synthetic peptides of the OmpB protein of *Rickettsia*, it is not possible to compare the results with diagnostic strategies using similar peptides; only a multi-peptide strategy with optimal results in sensitivity and specificity has been reported. Although the reactivity obtained is comparable to experimental ELISA techniques using pure antigens for *Rickettsia*, which achieve fluctuating sensitivity and specificity values between 80 and 100%.^11,13,30,31^

## 5. CONCLUSION

This study obtained by reverse vaccinology a set of peptides that were evaluated as antigens against previously diagnosed serum corresponding to *Rickettsia sp* and Dengue virus on nitrocellulose membranes, suggesting that peptides obtained by *in silico* methods may be useful for the development of accessible, simple, and affordable diagnosis for those diseases in endemic areas

## 6 ACKNOWLEDGEMENTS

Our thanks to the laboratory of emerging and reemerging diseases of the Dr. Hideyo Noguchi Regional Research Center for allowing the use of equipment, materials, and reagents. To the Institutional Postgraduate in Health Sciences by Universidad Autónoma de Yucatán and to CONAHCYT. This study was part of an MSc dissertation by Carlos Pena Bates supervised by Dra. Karla Rossanet Dzul-Rosado and Dr. Cesar Lugo Caballero.

## AUTHOR SUMMARY

**Carlos Peña**: Conceptualization, Methodology, Investigation, Writing-Original draft preparation. **Cesar Lugo**: Supervision, Conceptualization, methodology, Investigation, Writing-Reviewing and Editing, Resources. **Norma Pavía**: Resources. **Henry Noh**: Methodology, Investigation. **Karla Dzul**: Supervision, Conceptualization, methodology, Investigation, Writing-Reviewing and Editing, Resources. **Oghenekaro Omodior**: Writing-Reviewing and Editing. **Fernando Puerto**: Writing-Reviewing and Editing, Resources.

## 9. SUPPORTING INFORMATION

## S1

**Table S1.1.** Serums selected in the characterization of reference samples to evaluate Dengue virus peptides.

| Sera | Age<br>(Years) | Sex | *Rickettsia* |  | DENV2 |  |
| --- | --- | --- | --- | --- | --- | --- |
|  |  |  | *R. rickettsii* IgG/IgM | *R. typhi* IgG/IgM | ELISA IgG | Dot blot IgG |
| 1 | 15 | F | NEGATIVE | NEGATIVE | POSITIVE | NEGATIVE |
| 2 | 18 | F | NEGATIVE | NEGATIVE | POSITIVE | POSITIVE |
| 3 | 14 | F | NEGATIVE | NEGATIVE | POSITIVE | NEGATIVE |
| 4 | 22 | F | NEGATIVE | NEGATIVE | POSITIVE | POSITIVE |
| 5 | 54 | M | NEGATIVE | NEGATIVE | POSITIVE | POSITIVE |
| 6 | 8 | F | NEGATIVE | NEGATIVE | POSITIVE | POSITIVE |
| 7 | 30 | M | NEGATIVE | NEGATIVE | POSITIVE | POSITIVE |
| 8 | 11 | M | NEGATIVE | NEGATIVE | POSITIVE | POSITIVE |
| 9 | 54 | F | NEGATIVE | NEGATIVE | POSITIVE | NEGATIVE |
| 10 | 8 | M | NEGATIVE | NEGATIVE | POSITIVE | POSITIVE |
| 11 | 64 | M | NEGATIVE | NEGATIVE | POSITIVE | POSITIVE |
| 12 | 58 | M | NEGATIVE | NEGATIVE | POSITIVE | POSITIVE |
| 13 | 72 | M | NEGATIVE | NEGATIVE | POSITIVE | NEGATIVE |
| 14 | 36 | M | NEGATIVE | NEGATIVE | POSITIVE | POSITIVE |
| 15 | 20 | F | NEGATIVE | NEGATIVE | POSITIVE | POSITIVE |
| 16 | 39 | M | NEGATIVE | NEGATIVE | POSITIVE | NEGATIVE |
| 17 | 22 | F | NEGATIVE | NEGATIVE | POSITIVE | POSITIVE |
| 18 | 61 | M | NEGATIVE | NEGATIVE | POSITIVE | POSITIVE |
| 19 | 15 | F | NEGATIVE | NEGATIVE | POSITIVE | POSITIVE |
| 20 | 16 | F | NEGATIVE | NEGATIVE | POSITIVE | POSITIVE |

**Table S1.2.**
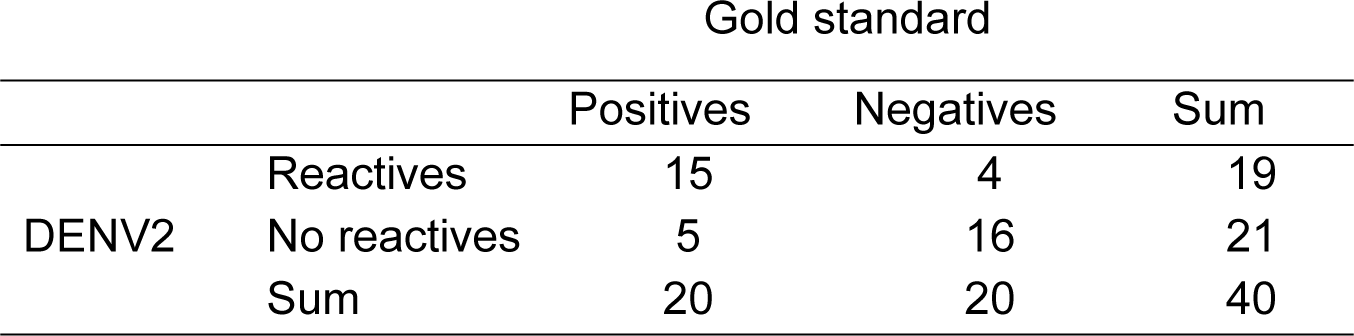
Contingency table comparing gold standard versus synthetic peptide dot blot.

## S2

**Table S2.3.** Serums were selected in the characterization of reference samples to evaluate Rickettsia peptides.

| Sera | Age<br>(years) | Sex | IFI *R. rickettsi* |  | IFI *R. typhi* |  | Dot blot |  |
| --- | --- | --- | --- | --- | --- | --- | --- | --- |
|  |  |  | Ig G | Ig M | Ig G | Ig M | IgG | IgM |
| 1 | 48 | F | 1: 256 | 1:128 | 1:128 | 1:128 | POSITIVE | POSITIVE |
| 2 | 16 | F | 1:128 | 1:32 | 1:128 | 1:64 | POSITIVE | POSITIVE |
| 3 | 33 | F | 1: 64 | NEGATIVE | 1:128 | NEGATIVE | POSITIVE | POSITIVE |
| 4 | 16 | F | 1:64 | NEGATIVE | 1:64 | 1:32 | POSITIVE | POSITIVE |
| 5 | 19 | F | 1:128 | NEGATIVE | 1:64 | NEGATIVE | NEGATIVE | NEGATIVE |
| 6 | 11 | M | 1:64 | 1:64 | 1:128 | 1:64 | POSITIVE | POSITIVE |
| 7 | 3 | F | NEGATIVE | NEGATIVE | 1:256 | 1:128 | POSITIVE | NEGATIVE |
| 8 | 3 | F | 1:128 | 1:64 | NEGATIVE | NEGATIVE | POSITIVE | POSITIVE |
| 9 | 2.5 | M | 1:64 | NEGATIVE | NEGATIVE | NEGATIVE | NEGATIVE | NEGATIVE |
| 10 | 57 | M | 1:128 | 1:256 | 1:128 | 1:128 | POSITIVE | POSITIVE |
| 11 | 3 | M | 1:64 | NEGATIVE | 1:128 | NEGATIVE | POSITIVE | NEGATIVE |
| 12 | 3 | M | 1:256 | NEGATIVE | 1:512 | NEGATIVE | POSITIVE | POSITIVE |
| 13 | 5 | F | NEGATIVE | 1:128 | 1:64 | NEGATIVE | POSITIVE | POSITIVE |
| 14 | 6 | F | 1:64 | NEGATIVE | 1:64 | 1:128 | POSITIVE | NEGATIVE |
| 15 | 4 | M | 1:64 | 1:64 | 1:64 | 1:128 | NEGATIVE | POSITIVE |
| 16 | 5 | M | NEGATIVE | 1:128 | NEGATIVE | 1:64 | POSITIVE | POSITIVE |
| 17 | 17 | F | 1:64 | 1:128 | 1:64 | 1:256 | POSITIVE | POSITIVE |
| 18 | 11 | M | 1:128 | 1:512 | 1:64 | 1:128 | POSITIVE | POSITIVE |
| 19 | 8 | F | 1:128 | 1:256 | 1:64 | 1:64 | POSITIVE | POSITIVE |
| 20 | 72 | M | 1:128 | 1:256 | 1:128 | 1:64 | POSITIVE | POSITIVE |

**Table S2.4.** Contingency table comparing gold standard versus synthetic peptide dot blot.

|  |  |  | Gold standard |  | Sum |
| --- | --- | --- | --- | --- | --- |
|  |  |  | Positives | Negatives |  |
| OmpB<br>peptides in<br>dot blot | IgG | Reactives | 17 | 3 | 20 |
|  |  | No reactives | 3 | 17 | 20 |
|  |  | Sum | 20 | 20 | 40 |
|  | IgM | Reactives | 15 | 4 | 19 |
|  |  | No reactives | 2 | 19 | 21 |
|  |  | Sum | 17 | 23 | 40 |

## S3. Flowchart of bioinformatic methods performed

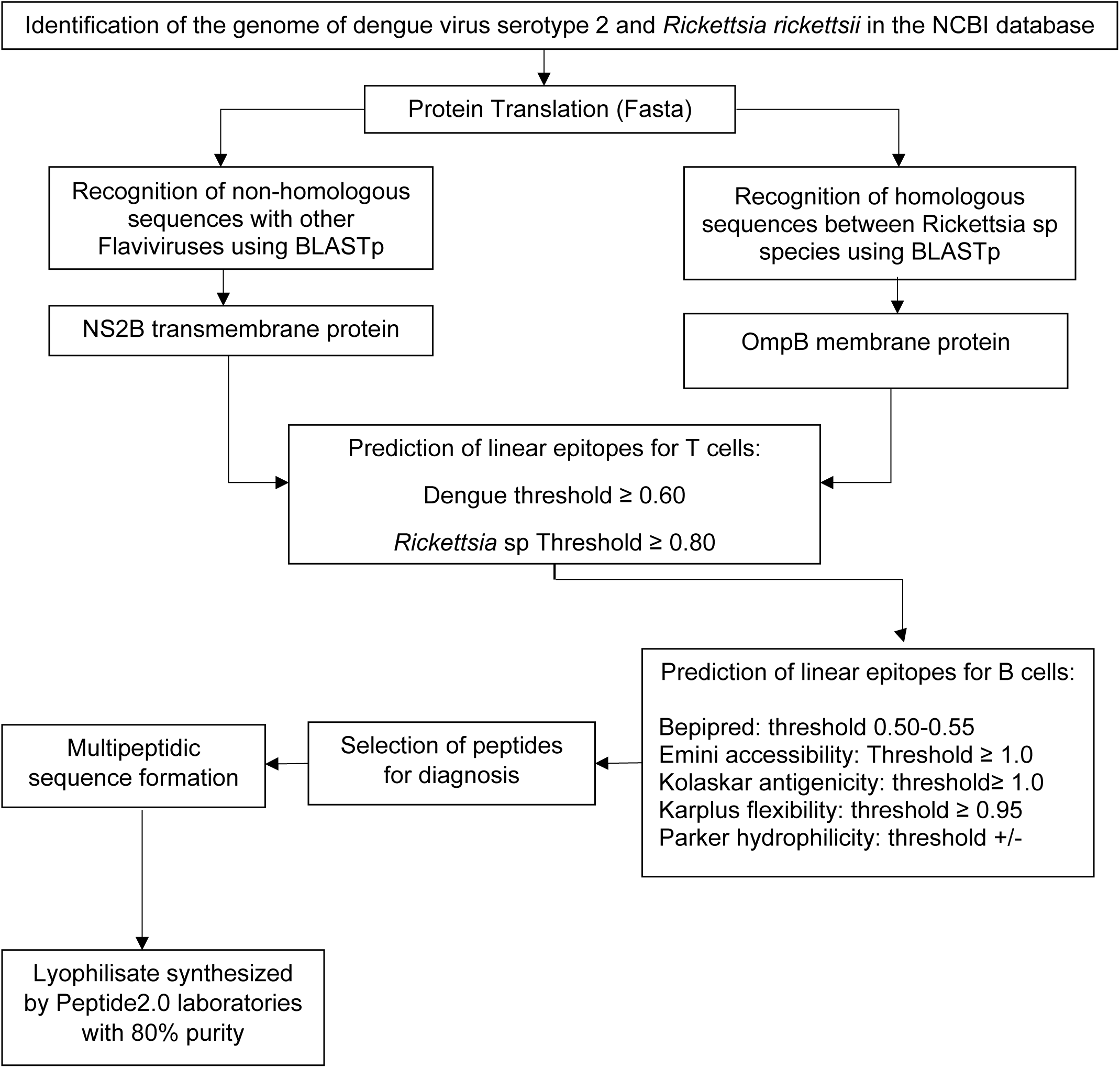

